# Identifying Changepoints in Biomarkers During the Preclinical Phase of AD

**DOI:** 10.1101/508820

**Authors:** Laurent Younes, Marilyn Albert, Abhay Moghekar, Anja Soldan, Corinne Pettigrew, Michael I Miller

**Affiliations:** Department of Applied Mathematics and Statistics, Johns Hopkins University; Department of Neurology, Johns Hopkins University; Department of Biomedical Engineering, Johns Hopkins University

**Author notes:** Correspondence: Laurent Younes, Department of Applied Mathematics and Statistics, Johns Hopkins University, 3400 N Charles st., Baltimore MD 21218. **Funding:** This study was supported in part by grants from the National Institutes of Health (U19-AG03365, P50-AG005146).

**Keywords:** Preclinical Alzheimer’s Disease, Biomarkers, Changepoints

## Abstract

**Objective:** Several models have been proposed for the evolution of Alzheimer’s disease (AD) biomarkers. The aim of this study was to identify changepoints in a range of biomarkers during the preclinical phase of AD.

**Methods:** We examined nine measures based on cerebrospinal fluid (CSF), magnetic resonance imaging (MRI) and cognitive testing, obtained from 306 cognitively normal individuals, a subset of whom subsequently progressed to the symptomatic phase of AD. A changepoint model was used to determine which of the measures had a significant change in slope in relation to clinical symptom onset.

**Results:** All nine measures had significant changepoints, all of which preceded symptom onset, however the timing of these changepoints varied considerably. A single measure, CSF-tau, had an early changepoint (40 years prior to symptom onset). A group of measures, including the remaining CSF measures (CSF-Abeta and phosphorylated tau) and all cognitive tests had changepoints 10-15 years prior to symptom onset. A second group is formed by medial temporal lobe shape composite measures, with a five-year time difference between the right and left side (respectively nine and three years prior to symptom onset).

**Conclusions:** These findings highlight the long period of time prior to symptom onset during which AD pathology is accumulating in the brain. There are several significant findings, including the early changes in cognition and the laterality of the MRI findings. Additional work is needed to clarify their significance.

## 1 Introduction

Accumulating evidence indicates that the underlying neuropathological mechanisms associated with Alzheimer’s disease (AD) begin a decade or more before the emergence of mild cognitive impairment (MCI) (Sperling, Aisen et al. 2011). This has led to an increasing interest in understanding the order and magnitude of biomarker changes during this ‘preclinical’ phase of AD.

A hypothetical model has been proposed describing the order in which biomarkers change across the spectrum of AD (Jack, Knopman et al. 2013). It has, however, been challenging to effectively test this model since most longitudinal studies that have enrolled cognitively normal individuals and collected relevant measures have limited follow-up. Additionally, studies with limited follow-up tend to lack a sufficient number of clinical outcomes (i.e., number of cases who progress to MCI) and therefore have limited power for statistical analyses designed to determine the timing of biomarker changes during preclinical AD.

Such analyses are feasible using data from the BIOCARD study, in which participants were cognitively normal when first enrolled, a wide range of informative measures were collected at baseline, and some participants have now been followed for over 20 years. The availability of these measures when the subjects were cognitively normal, and the unusually long duration of follow-up, allows the examination of the timing of biomarker changes during preclinical AD.

The primary goal of the analyses described here was to identify changepoints in measures based on cerebrospinal fluid (CSF), magnetic resonance imaging (MRI), and cognitive testing, obtained from a cohort of cognitively normal individuals, a subset of whom subsequently progressed to the symptomatic phase of AD. This study approved by the Johns Hopkins Medicine Institutional Review Board.

## 2 Methods

### 2.1 Study Design

The BIOCARD study, the study from which these data were drawn, was initiated at the National Institutes of Health (NIH) in 1995. While at the NIH, subjects were administered a neuropsychological battery and clinical assessments annually. MRI scans, CSF, and blood specimens were obtained approximately every two years. The study was stopped in 2005 for administrative reasons and re-established at Johns Hopkins University (JHU) in 2009, at which point the annual clinical and neuropsychological assessments were reinitiated. Bi-annual collection of CSF and MRI scans was re-established in 2015, and the acquisition of positron emission tomography (PET) scans using Pittsburgh Compound B (PiB) was begun. Tau PET imaging was initiated in 2017 (see Figure 1 for a schematic representation of the study design). This paper is based on CSF and MRI data collected during the 1995-2005 period and neuropsychological tests during the 1995-2013 period.

**Figure 1:**
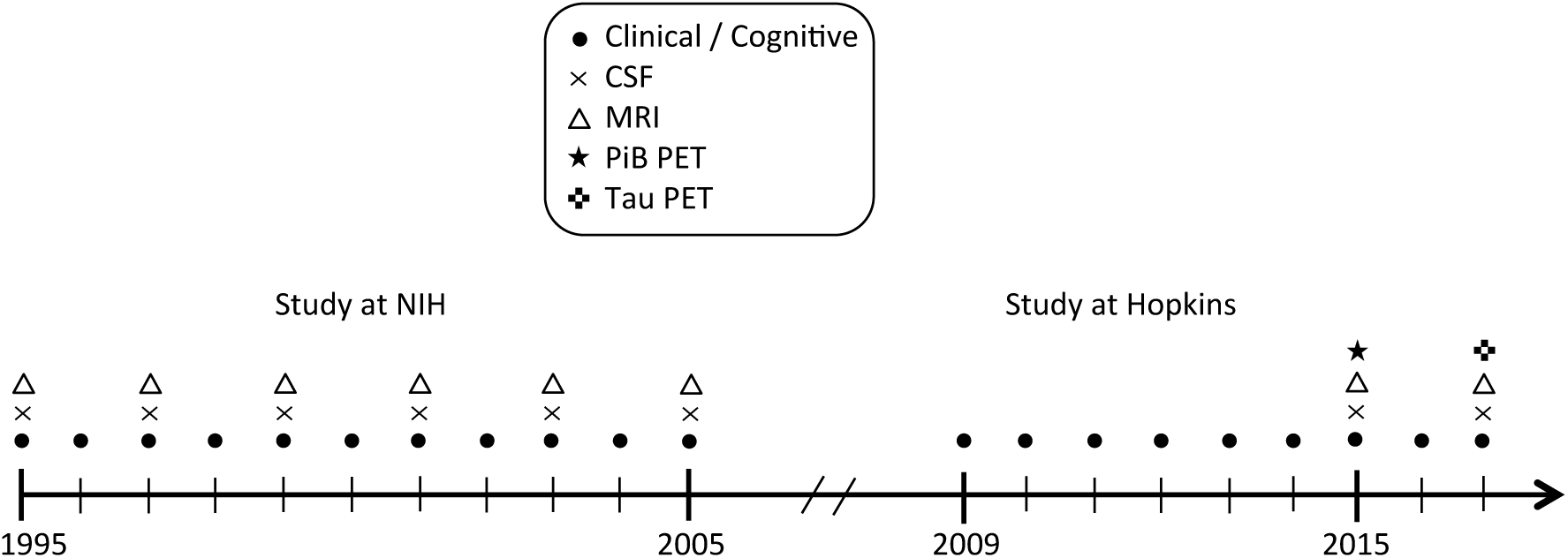
Schematic representation of the study design.

Qualified researchers may obtain access to all de-identified clinical and imaging data used for this study.

### 2.2 Selection of Participants

Recruitment was conducted by the staff of the Geriatric Psychiatry branch of the intramural program of the National Institute of Mental Health. At baseline, all participants completed a comprehensive evaluation at the NIH, consisting of a physical, neurological and psychiatric examination, an electrocardiogram, standard laboratory studies, and neuropsychological testing. Individuals were excluded from participation if they were cognitively impaired or had significant medical problems such as severe cerebrovascular disease, epilepsy or alcohol or drug abuse.

A total of 349 individuals were initially enrolled in the study, after providing written informed consent. By design, approximately 75% of the participants had a first degree relative with dementia of the Alzheimer type. The analyses presented here are based on data from 290 subjects who were cognitively normal at baseline and had complete observations on the baseline variables of interest. Subjects were excluded from analyses for the following reasons: (1) subjects had not yet re-enrolled in the study or had withdrawn (*n* = 29); (2) Subjects were below 40 years old at the beginning of study (*n* = 20). It should be noted that not all biomarkers were available for every subject and the actual number of subjects used for each run of the model varied: 256 for CSF, 270 for MRI and 281 for cognitive tests, for which we also excluded subjects who were only administered the test battery on one occasion, to allow for a more reliable practice effect correction (see Section 2.8).

Of the 290 subjects included in these analyses, 209 subjects remained cognitively normal at their last visit and 81 subjects were diagnosed with MCI or dementia due to AD by the time of their last visit. The demographic characteristics of the subjects in the analysis are shown in Table 1, which are similar to the characteristics of the cohort as a whole.

**Table 1:** Baseline Characteristics of the Participants Included in the Analyses in Comparison to the Cohort as a Whole.

| **Variable** | **Cohort as a whole** | **Subjects in Analysis** |
| --- | --- | --- |
| Subjects in Analysis | 349 | 290 |
| Age, mean years (SD) | 57.3 (10.4) | 58.8 (8.5) |
| Gender, % females | 57.6% | 58.3% |
| Education, mean years (SD) | 17.0 (2.4) | 17.1 (2.3) |
| Ethnicity, % Caucasians | 97.1% (0.9%) | 89.3% |
| % ApoE-4 carriers | 33.6% | 35.4% |
| MMSE, mean score (SD) | 29.5 (0.9) | 29.5 (0.8) |

Abbreviations: ApoE-4, apolipoprotein E-4; MMSE, Mini-Mental State Exam;

Abbreviations: ApoE-4, apolipoprotein E-4; MMSE, Mini-Mental State Exam;

### 2.3 Consensus Diagnostic Procedures

Clinical and cognitive assessments were completed annually at the NIH initially and subsequently at JHU, as noted above. A consensus diagnosis for each study visit was established by the staff of the BIOCARD Clinical Core at JHU (prospectively for subjects evaluated starting in 2009 and retrospectively for subjects evaluated at the NIH). This research team included: neurologists, neuropsychologists, research nurses and research assistants. During each study visit, each subject had received a comprehensive cognitive assessment and a Clinical Dementia Rating (CDR), as well as a comprehensive medical evaluation (including a medical, neurologic and psychiatric assessment). For the cases with evidence of clinical or cognitive dysfunction, a clinical summary was prepared that included information about demographics, family history of dementia, work history, past history of medical, psychiatric and neurologic disease, current medication use and results from the neurologic and psychiatric evaluation at the visit. The reports of clinical symptoms from the CDR interview with the subject and collateral source (e.g., spouse, child, friend) were summarized, and the results of the neuropsychological testing were reviewed.

The diagnostic process for each case was handled in a similar manner. Two sources of information were used to determine if the subject met clinical criteria for the syndromes of MCI or dementia: (1) the CDR interview conducted with the subject and the collateral source was used to determine if there was evidence that the subject was demonstrating changes in cognition in daily life, (2) cognitive tests scores (and their comparison to established norms) were used to determine if there was evidence of significant decline in cognitive performance over time. If a subject was deemed to be impaired, the decision about the likely etiology of the syndrome was based on the medical, neurologic, and psychiatric information collected at each visit, as well as medical records obtained from the subject, where necessary. More than one etiology could be endorsed for each subject (e.g., Alzheimer’s disease and vascular disease). One of four possible diagnostic categories was selected at each visit for each subject: (1) Normal, (2) Mild Cognitive Impairment, (3) Impaired Not MCI or (4) Dementia. The decision about the estimated age of onset of clinical symptoms was determined separately, and was based on responses from the subject and collateral source during the CDR interview regarding approximately when the relevant clinical symptoms began to develop. These diagnostic procedures are comparable to those implemented by the Alzheimer’s Disease Centers program supported by the National Institute on Aging.

The estimated age of onset of clinical symptoms was based primarily on a semi-structured interview with the subject and the collateral source. The staff conducting the consensus diagnoses were blinded to the CSF and imaging measures.

Within the context of this study, the diagnosis of Impaired Not MCI typically reflected contrasting information from the CDR interview and the cognitive test scores (i.e., the subject or collateral source expressed concerns about cognitive changes in daily life but the cognitive testing did not show changes, or vice versa, the test scores provided evidence for declines in cognition but neither the subject nor the collateral source reported changes in daily life).

### 2.4 Selection Criteria for Variables Included in the Analyses

The changepoint analyses presented here include variables from the three primary domains evaluated in the BIOCARD study, obtained when subjects were first enrolled. These domains include: (1) cognitive test scores, (2) CSF values, and (3) MRI measures. In order to be as parsimonious as possible, we based the selection of which specific variables should be included in the analyses on findings from prior publications (Albert, Zhu et al. 2018) that examined each of these measures in relation to time to onset of clinical symptoms. A total of 9 measures were included, as described below.

### 2.5 Cognitive Assessments

The annual, comprehensive neuropsychological battery covered all major cognitive domains, including memory, executive function, language, visuospatial ability, attention, speed of processing and psychomotor speed (see (Albert, Soldan et al. 2014) for the complete battery). We selected four cognitive measures to include in the changepoint analyses, as these four measures were significant in the multivariate Cox models examining the association between baseline performance and time to onset of clinical symptoms: (1) Digit Symbol Substitution Test from the Wechsler Adult Intelligence Scale – Revised; (2) Logical Memory – delayed recall from the Wechsler Memory Scale – Revised; (3) Verbal Paired Associates – Immediate recall from the Wechsler Memory Scale - Revised; and (4) Boston Naming Test.

### 2.6 CSF Assessments

The CSF specimens collected from the participants were analyzed using the xMAP-based AlzBio3 kit [Innogenetics] run on the Bioplex 200 system. CSF specimens were analyzed in triplicate on the same plate. The CSF specimens were analyzed using the AlzBio3 kit. The AlzBio3 kit contains monoclonal antibodies specific for Aβ1-42 (4D7A3), t-tau (AT120), and p-tau181p (AT270), each chemically bonded to unique sets of color-coded beads, and analyte-specific detector antibodies (HT7, 3D6). Calibration curves were produced for each biomarker using aqueous buffered solutions that contained the combination of the three biomarkers at concentrations ranging from 54 to 1,799 pg/ml for synthetic Aβ1-42 peptide, 25 to 1,555 pg/ml for recombinant tau, and 15 to 258 pg/ml for a tau synthetic peptide phosphorylated at the threonine 181 position (i.e., the p-tau181p standard). Each subject had all samples (run in triplicate) analyzed on the same plate. The intra-assay coefficients of variation (CV) for plates used in this study were: 7.7% +/-5.3 (Aβ_1-42_); 7.1% +/- 4.9 (tau); 6.3% +/- 4.8 (p-tau_181_). Interassay (plate-to-plate) CVs for a single CSF standard run on all plates used in this study were: 8.9% +/-6.5 (Aβ _1-42_); 4.7% +/-3.3 (tau), and 4.3% +/-3.18 (p-tau_181_). Compared with studies using the same kits and platforms, our absolute results are at the median levels for Aβ_1-42_, tau, and p-tau_181_. The CVs, plate-to-plate variability, and the dynamic range of our assays are well within published norms (Mattsson, Zetterberg et al. 2009, Shaw, Vanderstichele et al. 2009).

Three CSF variables were generated from these analyses: (1) Abeta 42, (2) total tau (t-tau) and (3) phosphorylated tau (p-tau)) (Moghekar, Li et al. 2013).

### 2.7 MRI Assessments

The MRI scans acquired from the participants were obtained using a standard multi-modal protocol with a GE 1.5T scanner. The coronal scans employed an SPGR (Spoiled Gradient Echo) sequence (TR = 24, TE = 2, FOV = 256 × 256, thickness/ gap = 2.0/0.0 mm, flip angle = 20, 124 slices). The scans were processed with a semi-automated method, using region-of-interest large deformation diffeomorphic metric mapping (ROI-LDDMM) techniques (Miller, Younes et al. 2013). More precisely, the MRI volumetric regions of interest (ROI) included the entorhinal cortex, hippocampus, and amygdala. For each of the three regions of interest, landmarks were placed manually in each MRI scan to mark the boundaries of the ROI, following previously published protocols (see (Miller, Younes et al. 2013), (Csernansky, Joshi et al. 1998) for the hippocampus, (Munn, Alexopoulos et al. 2007) for the amygdala and (Miller, Younes et al. 2013) for the entorhinal cortex). Next, a group template for the entorhinal cortex, hippocampus, and amygdala was created, based on the set of baseline MRI scans. The same set of landmarks was placed into this group template as in the individual subject scans. ROI-LDDMM procedures were then used to map the group template to the individual subject scans, using both landmark matching (Csernansky, Wang et al. 2000) and volume matching (Beg, Miller et al. 2005). The resulting segmented binary images for the entorhinal cortex, hippocampus and amygdala were used to calculate the volume of each structure, by hemisphere, by summing the number of voxels within the volume.

A medial temporal lobe composite was used in the present analyses, based on an average of the entorhinal cortex, hippocampus and amygdala. (Prior analyses showed that this composite is more strongly associated with CSF alterations that are an early marker of AD than the individual MRI measures taken separately (Gross, Hassenstab et al. 2017).) The measurements from the right and left hemisphere were examined separately.

The volumetric measurements of the entorhinal cortex, hippocampus and amygdala were normalized for head size by including total intracranial volume (ICV) as a covariate (Sanfilipo, Benedict et al. 2004). ICV was calculated using coronal SPGR scans in Freesurfer 5.1.0 (Segonne, Dale et al. 2004).

### 2.8 Statistical Analysis

#### 2.8.1 Overview

The overall goal of the changepoint analyses was to determine if each of the measures selected for analysis had a significant changepoint in relation to time to onset of clinical symptoms and, if so, the timing of these changepoints with respect to one another. The model used in these analyses has previously been applied to MRI data in this cohort in order to establish the order in which changes occur in the volume, thickness and shape of medial temporal lobe regions during preclinical AD.(Younes, Albert et al. 2014) A more advanced version of the model (Tang, Miller et al. 2017) is applied here to the full range of biomarkers available in the study.

The changepoint is represented in the model as a significant change in slope (see Figure 2). The model uses all of the available data (both from subjects who remained normal as well for those who progressed to MCI) in order to estimate the changepoint. The main features of the model are as follows.

**Figure 2:**
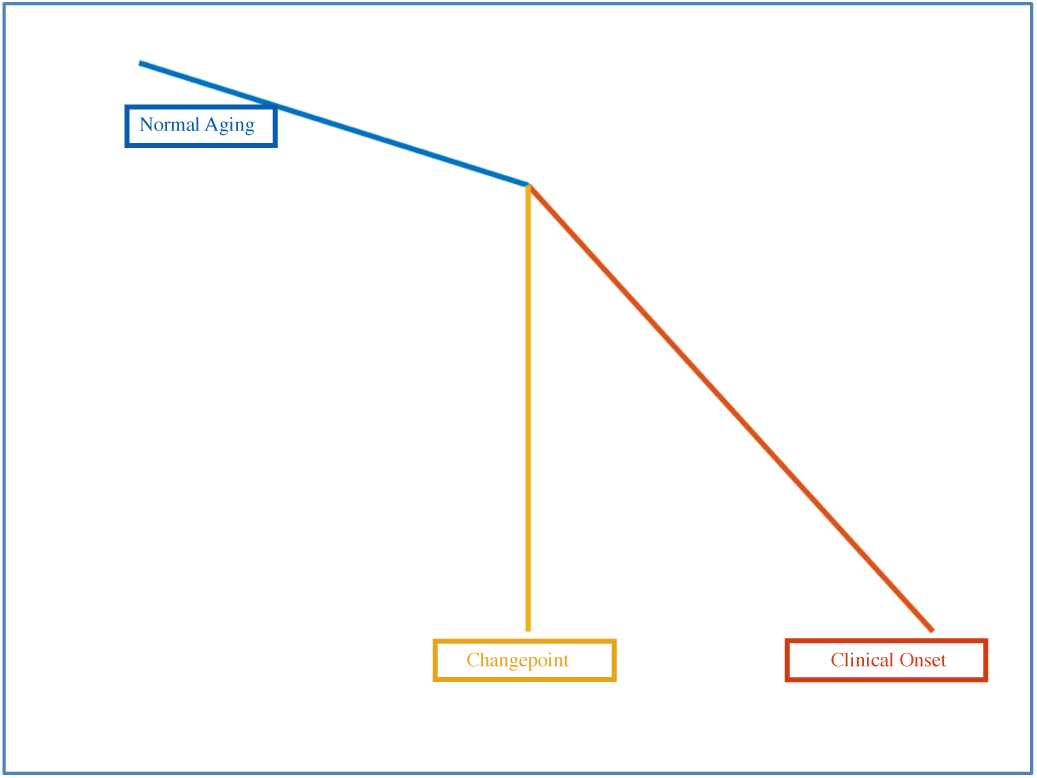
Schematic representation of the changepoint model.

- Time is measured relative to the clinical onset time of the disease (even though age is included as a covariate). This means that if a subject has been diagnosed with MCI ten years after another, the time scale for the latter is shifted ten years to the right compared with the former.
- Clinical onset times for normal subjects, which are not observed, are treated as missing data, therefore assuming “right censoring”. The model therefore assumes that every subject will ultimately get the disease if they were to live indefinitely. A prior model of disease onset is used in conjunction.
- The model assumes linearity in the measures as a function of age, with the change of slope at the changepoint. It is slightly different from the sigmoidal model of (Jack, Knopman et al. 2013), in which biomarkers smoothly transition from a low-abnormality plateau to a high-abnormality plateau in an S-shaped curve. Since the data in the study pertain to individuals who were cognitively normal at baseline and remained normal or who progressed from normal to MCI, the model assumes that the subjects are either in the low-abnormality range or in a transition phase, thus not requiring adaptation to an S-shaped function which might allow for two changepoints.
- All models included age and gender as covariates and a constant random effect. Education was included as an additional covariate for the cognitive measures; intracranial volume and left-handedness was included as an additional covariate for the MRI measures.
- We also corrected for the impact of a practice effect on cognitive tests (Rabbitt, Diggle et al. 2004, Zehnder, Bläsi et al. 2007, Vivot, Power et al. 2016) by introducing a covariate that depends on the number of tests taken in the past, defined by *z*_*practice*_ =1 − 2^−*k*^, when a test is taken for the *k*th time.
- For cognitive tests, we also limited our analyses to subjects that had at least two measurements over the course of the study. (This restriction was not applied to other biomarkers).
- CSF-tau and p-tau were transformed to logarithmic scale in the analyses.
- The model can project the changepoint forward in time as well as backward, sometimes allowing for a changepoint that precedes the initiation of data collection. Although the goal is to identify a changepoint preceding the onset of symptoms, this two-phase model allows for the changepoint and clinical onset of symptoms to coincide.

As shown in Figure 4, the model organizes the estimated time of symptom onset for the biomarker values along a broken line. The model fits the data so that the subjects with less abnormal values (e.g., higher test scores) tend to be on the left side of the curve and therefore to have a longer estimated time to clinical symptom onset.

**Figure 3:**
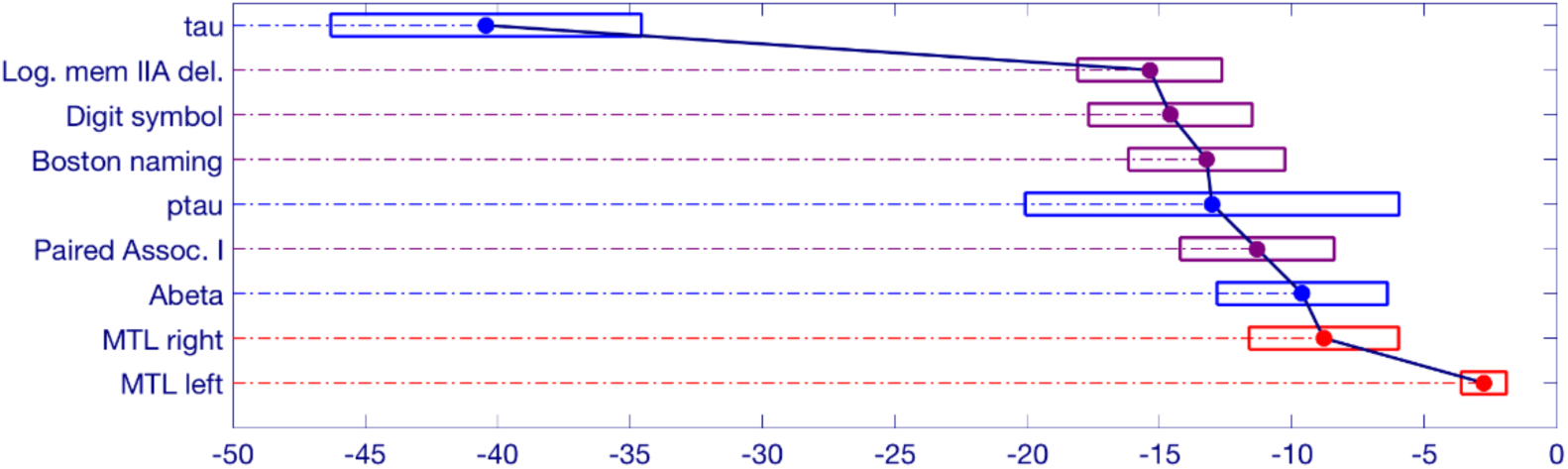
Schematic representation of significant changepoint results in relation to symptom onset. The estimated onset of clinical symptoms is represented by the value of 0 at the bottom right side of the figure. The numbers to the left of the 0 represent the estimated number of years prior to symptom onset for the changepoint of each variable. The width of each box represents a bias-corrected 75% confidence interval for the estimated value of each variable.

**Figure 4:**
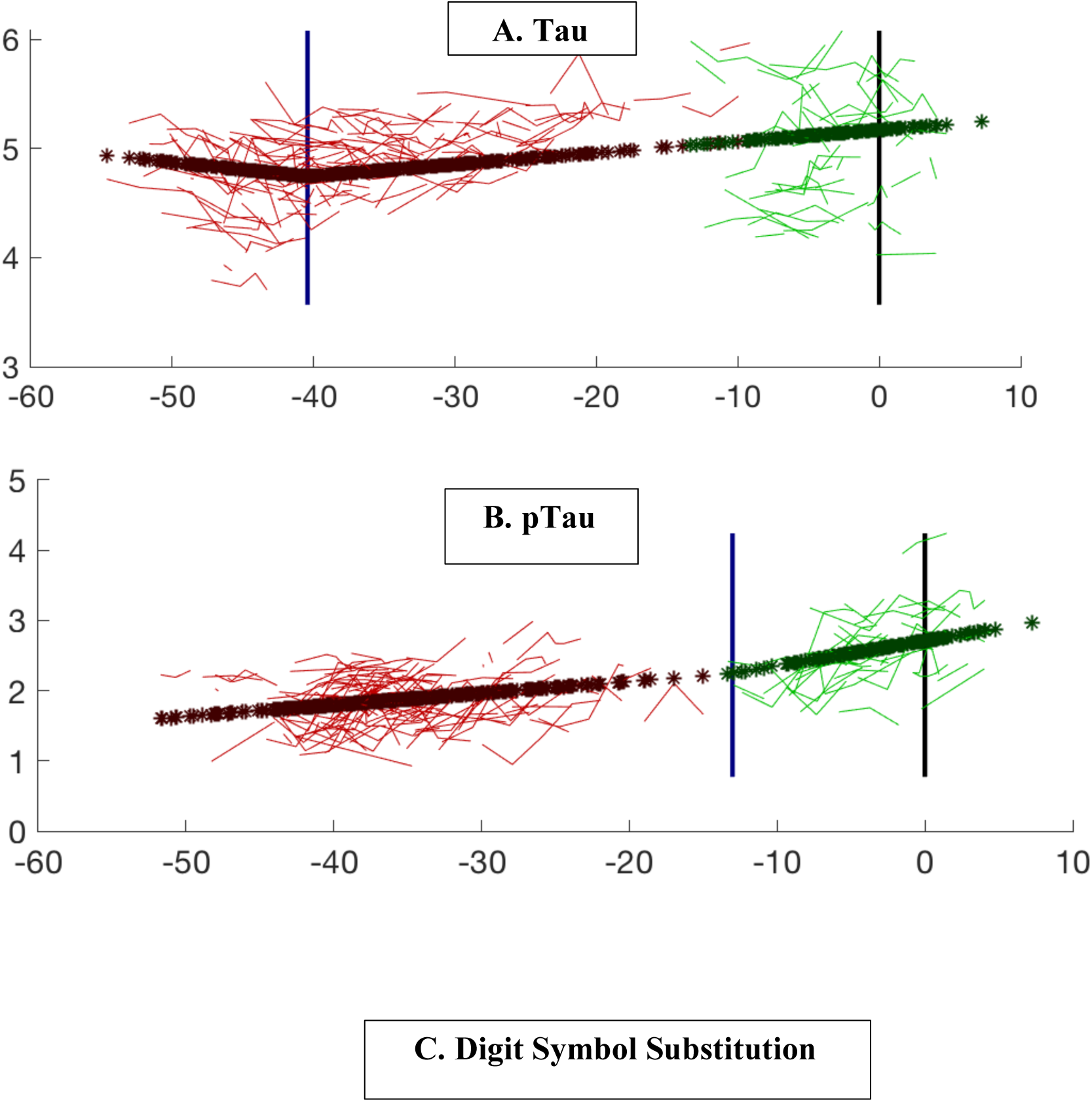
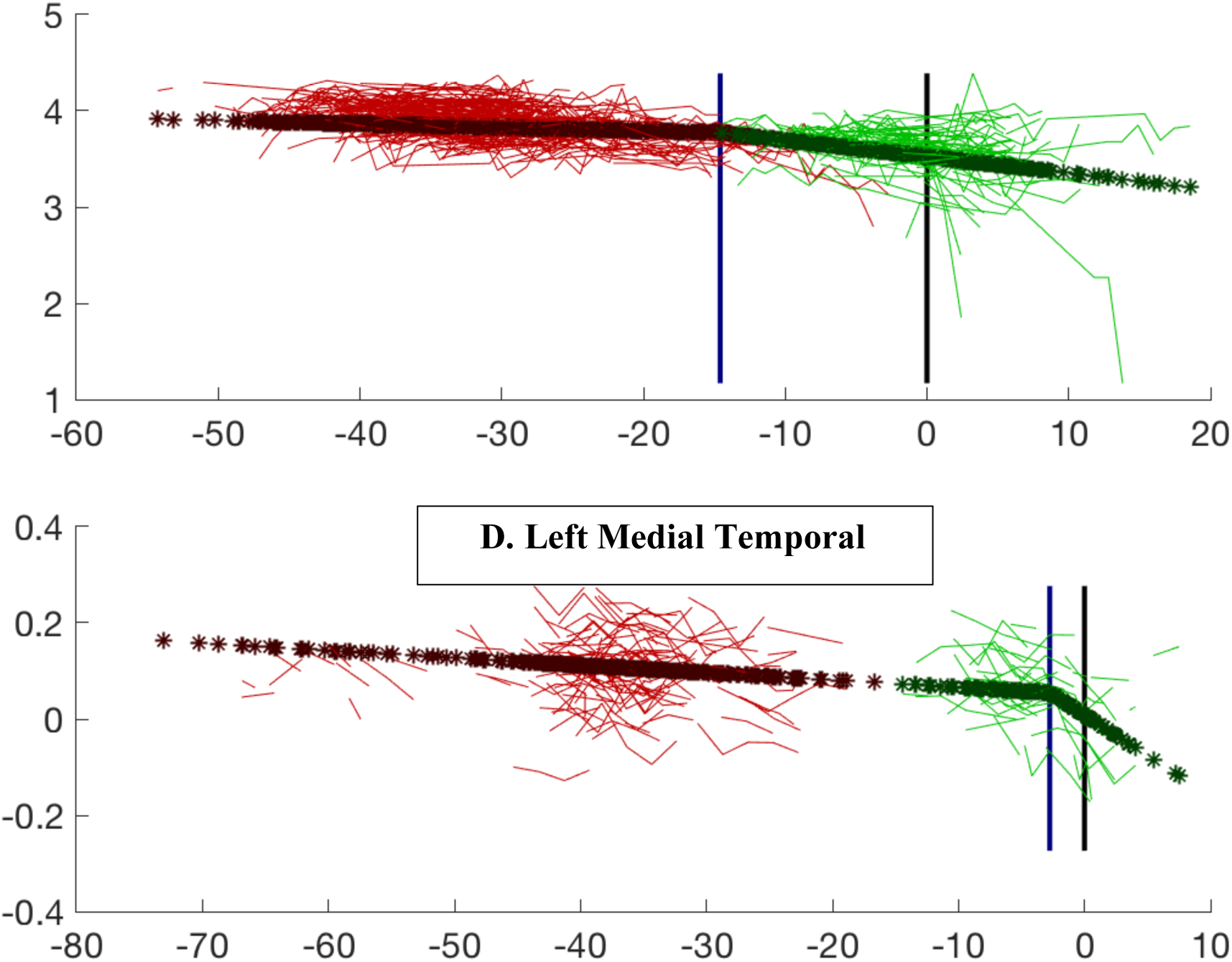
Model prediction for each variable compared with the observed data. The variables shown in the figures include: A. CSF tau; (B) CSF p-tau; (C) Digit Symbol Substitution Test; (D) Left Medial Temporal Lobe Volume. The red lines are the observed data for the subjects who remained cognitively normal. The green lines represent individuals who progressed to cognitive impairment. Dark red stars (and dark green stars, respectively) are the model predictions for the same subjects for whom observed data are presented. The blue vertical line marks the estimated changepoint. The black vertical line marks the estimated onset of clinical symptoms. The age of onset for the subjects who remained cognitively normal was imputed via Bayesian prediction.

#### 2.8.2 Mathematical description

Let *n* denote the number of subjects in the study. For the subject *k*, we assume *p*_*k*_ observations of a scalar biomarker, denoted 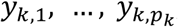, at ages 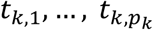. Let *T*_1_, …, *T*_*n*_ denote the subjects’ ages at the end of the study. Typically: 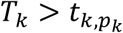 (age at last biomarker measurement). Let *U*_*k*_ denote the age at MCI onset, which is observed only if *U*_*k*_ ≤ *T*_*k*_.

Finally, let 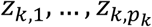 denote additional covariates, such as gender, education level, intracranial volume, etc. (Each *z*_*k*_ may be a vector.) Let η_*k*_ denote a constant random effect associated with each subject and 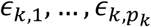 a random noise associated with each observation. They are modeled as Gaussian variables with respective variances τ^2^ and σ^2^. A prior distribution is used for *U*_*k*_, and modeled as a Gaussian with mean *m*_1_ = 93 years and standard deviation σ_1_ = 14.5 years. (This distribution was learned from a separate dataset.)

The changepoint model is

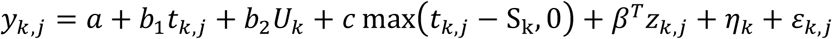

where *S*_*k*_ = max(*U*_*k*_ − Δ, 20) (in years) is the changepoint, the largest of Δ years before onset or 20 years.

This is a two-phase regression model. The biomarker first follows a linear trajectory (phase I)

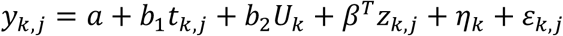

for *t*_*k,j*_ < *S*_*k*_ and then switches (with a continuous transition) to the model (phase II)

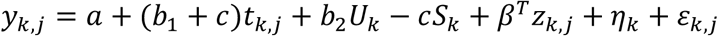

which is still linear, now with slope *b*_1_ + *c*.

The null hypothesis model assumes the phase I model over all times, or equivalently that *c* = 0.

The changepoint parameter is estimated using posterior means defined as follows. For each fixed Δ, the model parameters are estimated by maximum likelihood, and the value of the log-likelihood 𝓁(Δ) is computed. The estimator for Δ is then defined by

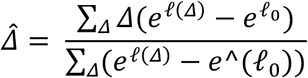

where the sum is over a finite number of Δ between 0 and 100, and 𝓁_0_ is the log-likelihood for the null hypothesis of no changepoint.

#### 2.8.3 Validation

A bootstrap method was used to evaluate significance and to assess the accuracy of the estimated onset time by providing confidence intervals of the estimated changepoint. This method, which estimates the standard errors based on random resampling with replacement, was used because it can be a more robust approach for calculating standard errors and statistical significance than parametric methods.

P-values and confidence intervals are estimated using bootstrap techniques. The bootstrap method estimates standard errors based on random resampling of the data with replacement; it can be a more reliable method of calculating standard errors and statistical significance than parametric methods (Efron 1979). For p-values, a general model is fitted, residuals are estimated, then resampled to reconstruct a model satisfying the null hypothesis (hence with *c* = 0). For confidence intervals, the approach is similar, but the full estimated model is used for reconstruction. We used 1,000 bootstrap samples for each estimation. A median absolute deviation was calculated for each changepoint in order to provide a robust estimate of the standard deviation, estimated as the median of the absolute value of the difference from the median of the sample as a whole.

Additionally, a ‘precedence graph’ was developed using the variables for which significant changepoints were calculated (n=8). Each of the measures were compared with one another using a bootstrap technique to determine the fraction of bootstrap samples for which the changepoint estimates for one measure were found to be earlier than the other. More precisely, precedence between two modalities A and B, with estimated changepoints 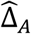 and 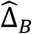 is assessed by computing the probability

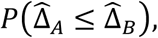

this probability being itself estimated using bootstrap resampling (i.e., for the sampling distribution). Because this probability requires to sample from the joint distribution of 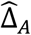 and 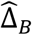, bootstrap samples are generated consistently across modalities, and the corresponding normalized frequency are computed over 1,000 replicas. This means that if, in order to reconstitute a bootstrap sample for visit time *t*′ in modality A, one has used the original residual computed at visit time *t*, then the corresponding original residual from the same visit time *t* in modality B will be used, whenever possible, to reconstitute a bootstrap sample for at time *t*′ for B. (When these two modalities have not been measured together at the considered visit times, the bootstrap samples are created independently.)

Groups of variables were computed using hierarchical clustering, based on precedence probability vectors, in order to provide a more concise representation of the changepoints with respect to one another. Arrows were drawn between measures when the confidence for one changepoint was earlier than the other at least 75% of the time.

#### 2.8.4 Null hypothesis model

The model with *c* = 0, which corresponds to our null hypothesis of no changepoint related to disease onset takes the form

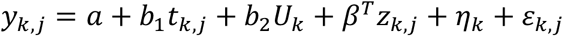

Importantly, it includes a disease effect, through the use of the onset time as a partially-observed covariate. While our primary focus here is on changepoint, there is certainly an interest in testing the significance of the hypothesis *b*_2_ ≠ 0, with respect to the “double-null” model

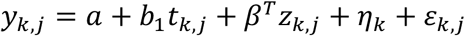

Which, this time, includes no disease related effect. A significant value of *b*_2_, say, with *b*_2_ > 0, implies a lower value of the biomarker for earlier cognitive onsets.

#### 2.8.5 Accounting for a normal changepoint

The changepoint in the proposed model is specified in terms of “time before disease onset” and does not include the possibility of such a change being due to normal aging. One of the difficulties in trying to account for both effects (let us call them disease vs. normal changepoint) is that if one of them is strong enough and not corrected for, it may induce significance when testing for the other effect even if that one is not present. On the other hand, correcting for an effect that is not present may reduce the power for detecting the other effect, even if the latter is present. For clarity and to simplify the exposition, we have focused our model and results on a single changepoint measured against disease onset. To be complete, however, we also explored a model in which a correction for a normal changepoint is included (which will therefore be more conservative for the detection of a change associated with disease). This model includes one additional covariate taking the form max (*t*_*kj*_ - δ, 0) where δ is a subject-independent age measuring the normal changepoint. To simplify the estimation process, this time δ is computed first (using maximum likelihood for a model without disease changepoint, which in this case only includes random effects as hidden variables), and then plugged into the general model.

## 3 Results

The results of the changepoint analyses for each of the nine variables examined are summarized in Table 2. As can be seen, all of the variables had a significant changepoint. The changepoints varied widely across the years preceding symptom onset (see Figures 3 and 4 for graphical representations of the model predictions).

**Table 2:** Results of Changepoint Analysis for CSF, MRI and Cognitive Variables

| **Variable Category/Name** | **Significance of Normal Changepoint p-value** | **Estimated Number of Years of Changepoint Prior to Symptom Onset [75% CI]** | **Median Absolute Deviation of Changepoint** |
| --- | --- | --- | --- |
| **Cerebrospinal fluid** |  |  |  |
| Abeta | < 0.001 | 9.6 [6.4-12.8] | 1.6 |
| p-tau | 0.024 | 13.0 [6.0-20.1] | 3.1 |
| tau | 0.002 | 40.4 [34.6-46.3] | 3.6 |
| **Magnetic Resonance Imaging** |  |  |  |
| Medial Temporal Lobe (L) | < 0.001 | 2.8 [1.9-3.6] | 0.2 |
| Medial Temporal Lobe (R) | 0.002 | 8.8 [6.0-11.6] | 1.5 |
| **Cognitive Test Scores** |  |  |  |
| Digit Symbol Substitution | < 0.001 | 14.6 [11.5-17.7] | 1.2 |
| Logical Memory Delayed | < 0.001 | 15.4 [12.6-18.1] | 1.6 |
| Paired Associates delayed | < 0.001 | 11.3 [8.4-14.2] | 1.2 |
| Boston Naming Test | 0.001 | 13.2 [10.3-16.2] | 1.7 |

The earliest changepoint is for CSF-tau, that is estimated at approximately 40 years. Significantly later in time, we estimate changepoints for two cognitive markers: The Logical Memory Delayed Recall (15.4 years prior to symptom onset), and the Digit Symbol Substitution Test (14.6 years prior to symptom onset). They are followed a couple of years later by the other two cognitive measurements: Boston Naming Test (13.2 years prior to clinical symptom onset) and Paired Associates Immediate Recall (11.3 years prior to symptom onset), with CSF p-tau in between (13.0 years prior to clinical symptom onset) and CSF abeta a little later (9.6 years prior to symptom onset). Imaging markers come next, with a six-year difference between the changepoints estimated on the right (8.8 years prior to clinical symptom onset) and on the left (2.8 years prior to clinical symptom onset) medial temporal lobe volumes. This arrangement is summarized in Figure 5 showing the precedence graph between these variables, in which arrows are placed only when the changepoint order could be estimated with enough reliability, as measured via bootstrap resampling.

**Figure 5:**
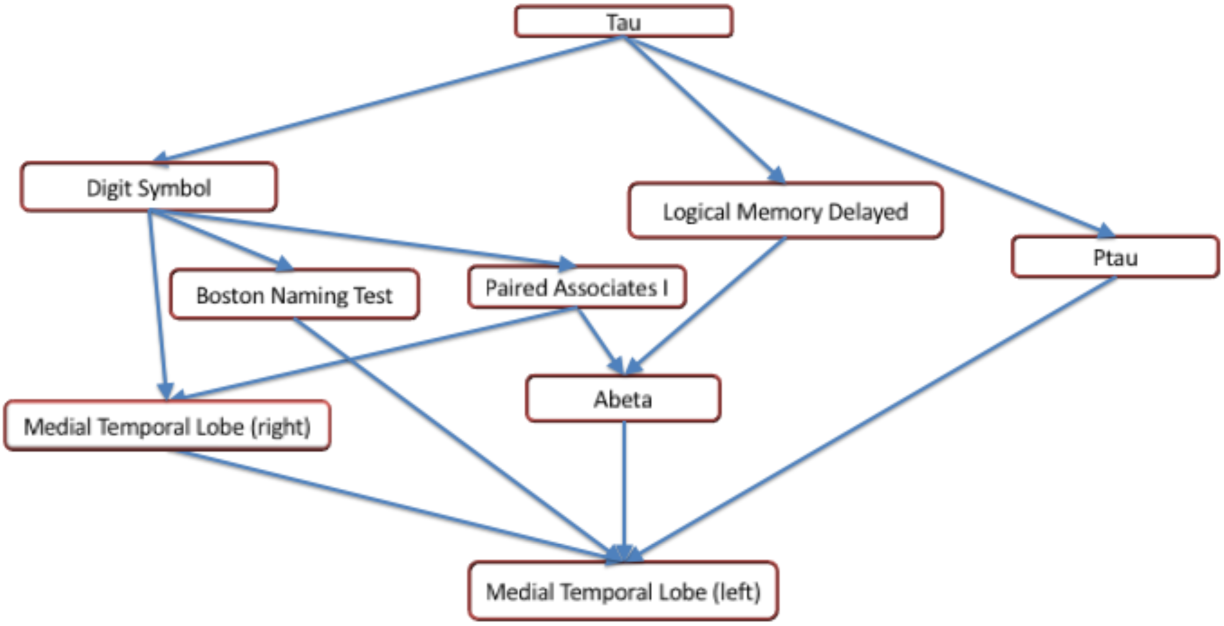
A precedence graph representing the order of the changepoints among the variables with significant changepoints. An arrow between groups of variables indicates that, more than 75% of the time, the changepoint for the variable represented as the ‘source’ was found to be earlier than the changepoint for the variable represented as the ‘target’, using bootstrap samples. The groupings of the variables were computed using hierarchical clustering within each modality, based on the precedence probability vectors. Arrows that can be inferred by transitivity are not shown for clarity.

All markers except CSF tau were significant for rejecting the double-null hypothesis of no effect of the cognitive onset time on the marker, with p-values given by 0.047 (left MTL), 0.004 (right MTL), 0.016 (CSF abeta), 0.007 (CSF p-tau) and less than 0.001 for all cognitive markers. We found *b*_2_ > 0 for all markers (indicating a smaller value of the marker for earlier onsets), except for CSF p-tau, for which this value is negative.

Variables that were significant for normal changepoints (p-value less than 0.05) at fixed age in the considered biomarkers for CSF p-tau (age: 51.3), CSF tau (age: 64.2 years), Digit Symbol Substitution (age: 74.8), Logical Memory Delayed (age: 67) and right medial temporal lobe (age: 48.6 years). Introducing this normal changepoint in the model as an additional covariate had limited impact on the significance and value of the disease changepoint times.

## 4 Discussion

The changepoint analyses presented here lead to several conclusions. First, the changepoint for CST tau occurs several decades prior to the onset of clinical symptoms. Second, the changepoints seen in the rest of the variables appear to reflect a cascade of events in which multiple measures are changing a decade prior to the onset of clinical symptoms. Third, there is a significant difference in the vulnerability of the right vs. the left medial temporal lobe. Several of these findings diverge from the hypothesized ordering of biomarkers in the model proposed by Jack et al (Jack, Knopman et al. 2013). as well as the hypothetical stages proposed in the NIA/AA Working Group Report, both of which propose that cognitive change follows significant accumulation of amyloid and tau (Sperling, Aisen et al. 2011). Our results describe a more complex ordering, in which some cognitive effects were found to predate changepoints in CSF abeta. It is of course possible that changes in amyloid occur at a time too early to be detectable in our model, or with a different slope associated with the disease, which is not addressed here.

To gain further insights into these findings, we looked in greater detail at the two cognitive tests with very early timepoints. Figure 4a (which shows the model regression on Tau after removal of covariate and random effects) indicates that, after a first phase during which Tau is almost flat, the protein appears to accumulate starting about 40 years before onset. This is a large gap, but, as already remarked, these results do not inform us on the pathological nature of this increase, but rather on the fact that an event/changepoint seems to happen for CSF-tau accumulation with a timing that can be associated with clinical impairment several decades later. While this changepoint appears to be associated with the onset of disease, it does not necessarily correspond to an early effect.

The difference in the changepoint for the right and left medial temporal has been presaged by prior reports that have examined the individual regions within the MTL separately. For example, we previously reported that both the right entorhinal cortex and amygdala, when measured at baseline, were significantly related to time to onset of symptoms, whereas measures on the left were not (Soldan, Pettigrew et al. 2015). Further studies are needed to determine why this differential vulnerability may occur.

It is important to acknowledge the limitations of these analyses. First, the wide confidence intervals for the CSF assays, particularly for CSF tau and p-tau, limit the ability to narrow down the changepoint for this important biomarker. The variability of CSF assays has been an acknowledged challenge in the field for some time, reflected by international efforts to develop improved methods (Mattsson, Andreasson et al. 2011). Newer assays are currently under development (Chang, Shan et al. July 2017) raising the possibility that measures with less variability will be soon available that will permit more accurate changepoint estimates, with narrower confidence intervals. Second, while the sample size used here is sufficient to generate findings with substantial statistical significance, the width of most of the 75% confidence intervals is between 5 and 10 years, some of them being even greater. Under the assumption that an increase in sample size may reduce the confidence intervals, we have established a consortium of five sites around the world that are collecting comparable data (Gross, Hassenstab et al. 2017). We plan to apply this changepoint model to data gathered from across the sites, which will greatly increase the sample size. Third, the model itself incorporates assumptions that limit its applicability. For example, a two-phase linear model assumes some continuity of the biomarkers before and after changepoint, since it only accounts for a change of slope. A very abrupt change, for example, would be imperfectly approximated by the model and may result in a loss of power in the likelihood ratio test. Sublinear or hyperlinear evolutions before or after changepoints may have a similar effect. There could also be more than one changepoint, which is not handled by the analysis thus far. Additionally, the estimates of the changepoint are generally more stable when the likelihood ratio test p-value is small, and for small changepoints. Lastly, the changepoint analysis presented here is for univariate biomarkers, and therefore it has been applied separately to each of the variables. While this approach can, in theory, be extended to the multivariate case, such an extension presents statistical challenges, which are currently under investigation.

In summary, in interpreting our results, significant results for our model provide a credible indication that a changepoint in the biomarker occurs some number of years before clinical onset. One of the two interpretations is that the changepoint marks an effect of the disease, which happens before its onset can be detected. A second interpretation, that we cannot rule out by the measurements we have made and tested, is that the change is non-pathological, but that its timing is correlated with the disease onset.

## Acknowledgements

The BIOCARD Study consists of 7 Cores with the following members: (1) the Administrative Core (Marilyn Albert, Barbara Rodzon, Corinne Pettigrew, Rostislav Brichko); (2) the Clinical Core (Marilyn Albert, Anja Soldan, Rebecca Gottesman, Ned Sacktor, Scott Turner, Leonie Farrington, Maura Grega, Gay Rudow, Daniel D’Agostino, Scott Rudow); (3) the Imaging Core (Michael Miller, Susumu Mori, Laurent Younes, Tilak Ratnanather, Timothy Brown, Anthony Kolasny, Kenichi Oishi, Andreia Faria); (4) the Biospecimen Core (Abhay Moghekar, Akhilesh Pandey, Jacqueline Darrow); (5) the Informatics Core (Roberta Scherer, Ann Ervin, Jennifer Jones, Hamadou Coulibaly, April Broadnax, David Shade); (6) the Biostatistics Core (Mei-Cheng Wang, Qing Cai, Jiangxia Wang); and (7) the Neuropathology Core (Juan Troncoso, David Nauen, Olga Pletnikova, Gay Rudow, and Karen Fisher).

The authors are grateful to the members of the BIOCARD Scientific Advisory Board who provide continued oversight and guidance regarding the conduct of the study including: Drs. John Csernansky, David Holtzman, David Knopman, Walter Kukull, and Kevin Grimm, and Drs. Laurie Ryan and John Hsiao, who provide oversight on behalf of the National Institute on Aging. The authors thank the members of the BIOCARD Resource Allocation Committee who provide guidance regarding the use of the biospecimens collected as part of the study, including: Drs. Constantine Lyketsos, Carlos Pardo, Gerard Schellenberg, Leslie Shaw, Madhav Thambisetty, and John Trojanowski.

The authors acknowledge the contributions of the Geriatric Psychiatry Branch of the intramural program of NIMH who initiated the study (Principal investigator: Dr. Trey Sunderland). The authors are particularly indebted to Dr. Karen Putnam, who has provided documentation of the Geriatric Psychiatry Branch study procedures and the data files received from the National Institute of Mental Health.

Author Contributions
Dr. Younes conducted the analyses and drafted the manuscript
Dr. Moghekar played a major role in acquiring the data
Dr. Albert interpreted the data and revised the manuscript
Dr. Soldan revised the manuscript for intellectual content
Dr. Pettigrew revised the manuscript for intellectual content
Dr. Miller interpreted the data and revised the manuscript

Disclosures
Dr. Younes reports no disclosures
Dr. Moghekar reports no disclosures
Dr. Albert is a consultant to Eli Lilly
Dr. Soldan reports no disclosures
Dr. Pettigrew reports no disclosures
Dr. Miller reports he has significant ownership in Anatomy Works, LLC, a relationship which is being handled by the University.

